# Persistent anti-heart autoimmunity causes cardiomyocyte damage in chronic heart failure

**DOI:** 10.1101/542597

**Authors:** Amalia Sintou, Sarah el Rifai, Catherine Mansfield, Stephen M. Rothery, Jose L. Sanchez Alonso, Salomon Narodden, Keshav Sharma, Elisa Ferraro, Muneer G. Hasham, Pamela Swiatlowska, Sian E. Harding, Nadia Rosenthal, Julia Gorelik, Susanne Sattler

## Abstract

Although clinicians and researchers have long appreciated the detrimental effects of excessive acute inflammation after myocardial infarction (MI), less is known about the role of the adaptive immune system in MI complications including heart failure. Yet, abundant cardiac self-antigens released from necrotic cardiomyocytes in a highly inflammatory environment are likely to overwhelm peripheral mechanisms of immunological self-tolerance and adaptive auto-reactivity against the heart may cause ongoing tissue destruction and exacerbate progression to chronic heart failure (CHF).

Here, we confirm that the adaptive immune system is indeed persistently active in CHF due to ischemic heart disease triggered by MI in rats. Heart draining mediastinal lymph nodes contain active secondary follicles with mature class-switched IgG2a positive cells, and mature anti-heart auto-antibodies binding to cardiac epitopes are still present in serum as late as 16 weeks after MI. When applied to healthy cardiomyocytes *in vitro,* humoral factors present in CHF serum promoted apoptosis, cytotoxicity and signs of hypertrophy.

These findings directly implicate post-MI autoimmunity as an integral feature of CHF progression, constituting a roadblock to effective regeneration and a promising target for therapeutic intervention.

---

Chronic heart failure (CHF) describes a pathological condition in which the heart is unable to pump enough blood to meet the metabolic demands of the body. It is the final outcome for a range of conditions including ischemic heart disease following myocardial infarction (MI) (1), and a major public health problem affecting 26 million people worldwide (2).

The immune system has been implicated in the early acute immune response to MI, which is crucial for quick tissue repair (3,4). Neutrophils and monocytes/macrophages infiltrating the damaged myocardium in response to danger signals (DAMPs) are an overt sign of early inflammation. However, the post-MI immune response does not cease once acute inflammation has been resolved, and excessive activation of endogenous repair mechanisms may lead to ongoing inflammation, fibrosis, and sustained tissue damage. Cardiac self-antigens released from necrotic cardiomyocytes in a highly inflammatory environment also activate autoreactive B and T cells of the adaptive immune system, which can be long lived and may cause sustained damage to the myocardium (4)(5).

The presence of anti-cardiac auto-antibodies in post-MI serum is well known to clinicians (6) (7). Post-myocardial infarction syndrome was first described in 1956 by Dressler (8) and is now recognized as an adaptive immune response against myocardial proteins (9). However, besides rare cases of clinically obvious extreme manifestations of post-MI autoimmunity, a certain degree of persistent subclinical auto-reactivity can cause ongoing damage to previously healthy myocardial tissue. This may increase susceptibility to subsequent infarcts and development of adverse remodeling and heart failure.

Activated B cells generate tissue-specific auto-antibodies and pro-inflammatory cytokines, which can directly contribute to cardiac dysfunction (37). B cell activation occurs in lymph nodes draining damaged tissue sites. Activated B and T cells with the same antigen specificity interact at the B-T border zone in the lymph node which leads to germinal centers facilitating affinity maturation and class switching to mature IgG isotypes, which convey a range of effector functions (10). Detailed information on the autoantibody isotypes in post-MI and CHF patients is limited, and further research may yield fundamental insights into pathological effects of auto-antibodies present in these patients.

In the present study, we document a highly active adaptive immune system in injured hearts, with anti-cardiac auto-reactivity and humoral factor-mediated cytotoxicity at chronic heart failure stage in the rodent MI model of permanent left anterior descending (LAD) artery ligation. We characterize the serum antibody isotype composition induced by myocardial injury and show a striking degree of immunological activity long after the initial injury. CHF serum creates a cytotoxic and hypertrophic environment for cultured cardiomyocytes, confirming the emerging notion that ongoing immune-mediated damage in the post-MI environment may promote heart failure, and constituting a new therapeutic focus for regenerative intervention.

## RESULTS

### 1) Persistent reactivity of heart-draining lymph nodes during CHF

An MI is a potent trigger for an acute local innate immune reaction, including early infiltration of neutrophils and macrophages (11)(11), but evidence is accumulating that T and B cells are also activated within a week post-MI (12). To investigate the activation state of the adaptive immune system in CHF progression, myocardial infarcts were induced in male Sprague Dawley rats by surgical ligation of the left anterior descending (LAD) coronary artery. Heart-draining mediastinal lymph nodes were isolated at week 2 and week 16 after infarct surgery and analyzed for signs of reactivity. They showed distinct lymphoid follicles containing active germinal centers acutely post-MI (2 weeks) as well as at chronic stage (16 weeks) post-MI. Blood was visible in the medullary sinuses of the week 2 post-MI lymph nodes, indicating that they drained an area of hemorrhage (Figure 1A), which is in line with the myocardial damage induced by LAD. Quantification of lymph node size, total number of follicles and the percentage of follicles with germinal centers confirmed an activated state at week 16, albeit decreased compared to week 2 (Figure 1B). Transcription of genes involved in B cell development and activation, including *Rag1, Cd20(MS4a1), Cd81* was upregulated in the spleen 2 weeks post-MI, and some were still elevated over baseline at CHF stage (Figure 1C). Recombination activating gene (*Rag1*) was elevated most prominently in both week 2 and week 16, consistent with its role in B cell development and Ig formation. *Rag* expression is also upregulated in antigen-activated early memory B cells during autoimmune responses (13). Sustained upregulation of *Il4* and the Th2-associated chemokine receptor *Ccr3* together with an increase in *Cd40* (*Tnfrsf5*) but a downregulation of *Cd40lg* (*Tnfsf5*) may provide an environment blocking apoptotic cell death and inducing sustained growth and differentiation (14).

**Figure 1:**
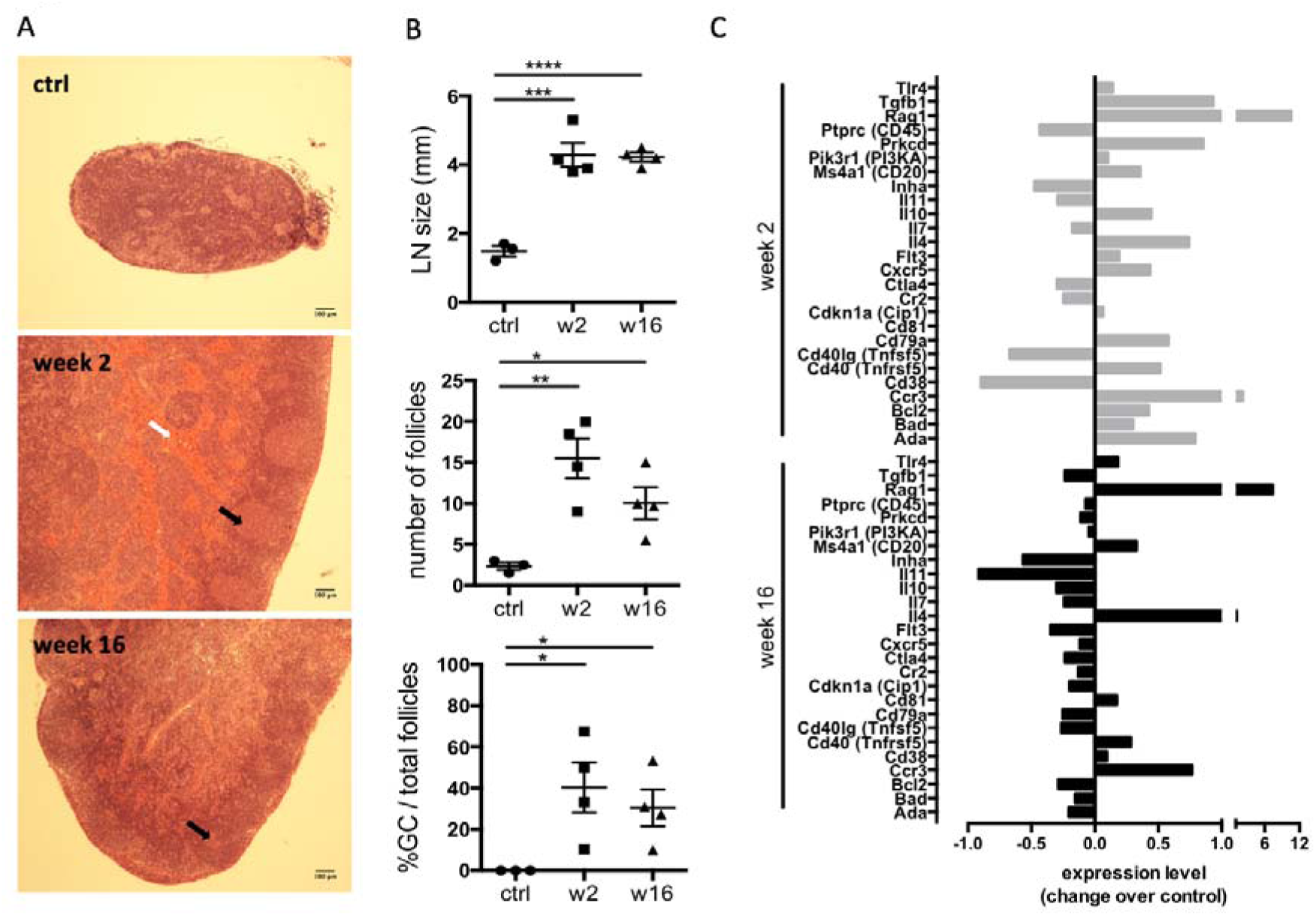
Persistent activation of heart-draining lymph nodes at CHF stage after infarct. Myocardial infarction was induced by surgical ligation of the LAD. Mediastinal (heart draining) lymph nodes were obtained from healthy rats (ctrl), 2 weeks (acute post-MI) and 16 weeks (CHF stage) post-MI. A: Histology (H&E) sections of representative lymph nodes. Post-MI rats show distinct lymphoid follicles containing active germinal centers (black arrows). Hemorrhage (grey arrow) is present in the medullary sinuses of the week 2 post-MI lymph nodes, indicating that they drain an area of hemorrhage. B: Lymph node size and quantification of the total number of follicles and the percent of follicles with germinal centers. C: qPCR array testing a range of genes involved in B cell activation. Bars represent expression change over baseline, with healthy baseline values set to 0. Experiment performed in triplicates on 3 pooled spleens of ctrl (healthy baseline), week 2 and week 16 post-MI rats. Statistics: n = 3 (ctrl) and 4 (week 2, 16), Values represent mean +/- s.e.m., Statistics: two-tailed Student’s t-test with Welch correction. *P<0.05, **P<0.005, ***P<0.001, ****P<0.0001.

### 2) The CHF antibody repertoire shifts towards mature class-switched isotypes

The presence of active germinal centers in heart-draining lymph nodes during CHF marks ongoing B cell maturation processes. To assess maturity of the CHF antibody repertoire, we characterized the isotype composition of the local and systemic antibodies in response to MI and their potential for pathological effects. Serum was collected 2 and 16 weeks post-MI and analyzed by ELISA for relative concentrations of immunoglobulin (Ig) light chains (IgKappa and IgLambda) and heavy chains (immature IgM, mature class-switched IgG1, IgG2a, b and c). A dramatic increase in overall antibody levels was evident (Figure 2a). While overall IgM-levels remained comparable to control (Figure 2b), mature IgG isotypes were increased, most prominently IgG2a (Figure 2c). Most strikingly, this was accompanied by a marked increase of IgG2a^+^ cells in the mediastinal lymph nodes, indicating heart specificity of mature class-switched autoreactive B cells (Figure 2d).

**Figure 2:**
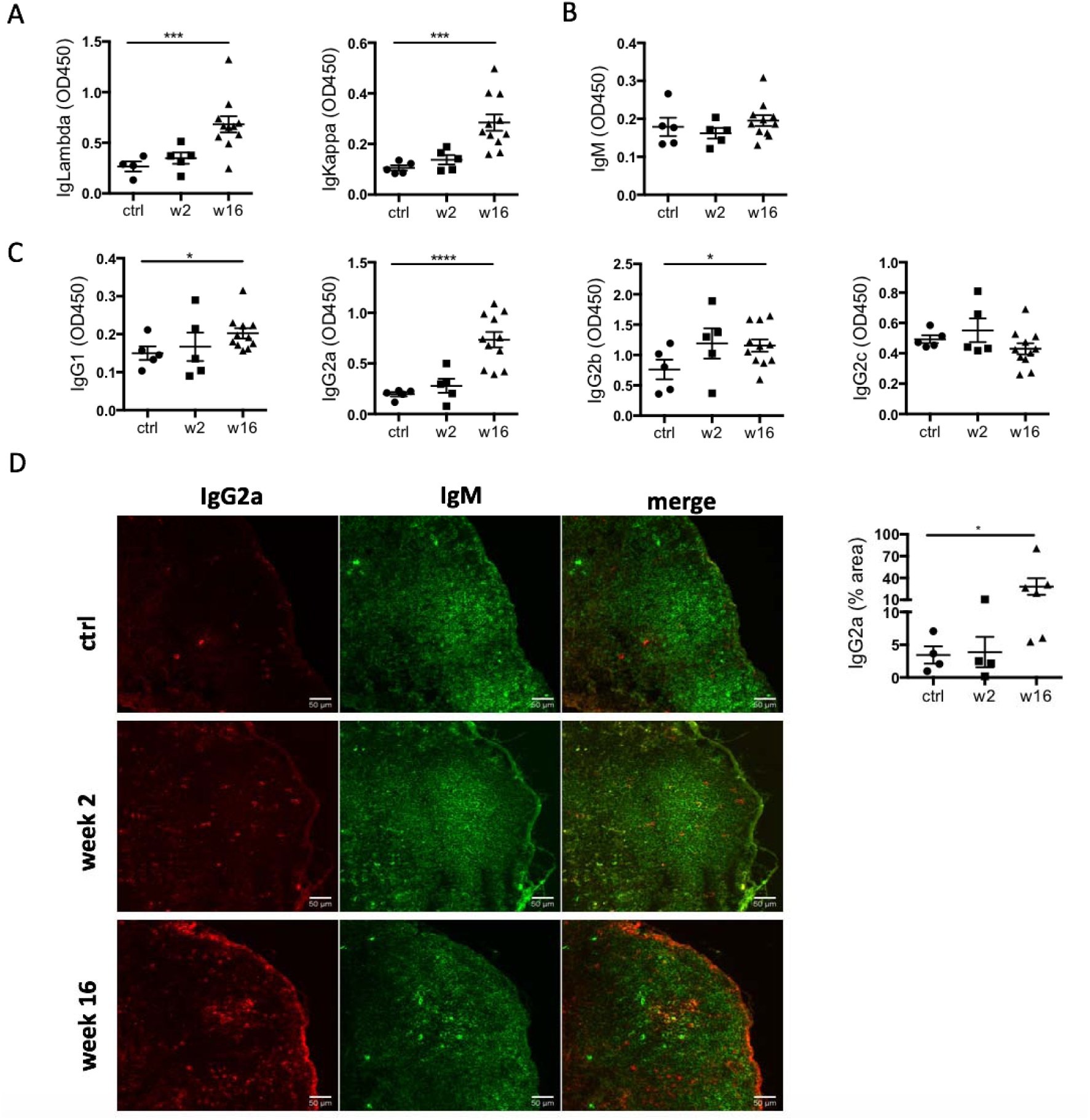
Levels of IgG2a antibodies in the circulation and IgG2a+ cells in the heart-draining lymph nodes are elevated during CHF. Myocardial infarction was induced by surgical ligation of the LAD. Serum was collected 2 weeks (acute post-MI) and 16 weeks (chronic heart failure stage) post-MI and analyzed by ELISA for relative concentration of A: immunoglobulin (Ig) light chains (IgKappa and IgLambda) and B, C: heavy chains (immature IgM (B), and mature class-switched IgG1, IgG2a, b and c (C). n = 5 (ctrl, week 2) and 11 (week 16) / group D: Immunofluorescence staining of mediastinal lymph nodes with Alexa Fluor®488 Goat anti-rat IgM (green) and Alexa Fluor®647 anti-rat IgG2a (red). n = 4 (ctrl, week 2) and 6 (week 16) / group, Statistics: values represent mean +/- s.e.m., two-tailed unpaired Student’s t-test with Welch correction. *P<0.05, ***P<0.001, ****P<0.0001.

### 3) CHF serum contains mature anti-heart auto-reactive antibodies

While the degree of overall increase in total antibody concentration and elevation of potentially pathological isotypes such as IgG2a in the periphery and the draining lymph nodes is striking in itself, CHF serum also contained auto-antibodies specific to cardiac protein. An ELISA assay using post-MI serum against coated healthy heart lysate showed a progressive increase in the levels of anti-heart IgG auto-antibodies (Figure 3a). Characterization of heart-specific auto-antibody population confirmed that it also contains mature class-switched IgG1, IgG2a and IgG2b isotype antibodies (Figure 3b). Although individuals show variable levels of IgG2a and IgG2b anti-heart auto-antibodies at CHF stage, elevation over baseline is evident in all animals. Notably, inbred Lewis rats of a different genetic background appeared protected from developing anti-heart auto-antibody levels supporting a difference in genetic susceptibility to post-MI autoimmunity (Supplementary Figure 1), most likely due to MHC haplotypes and antigen-depended mechanisms of B cell maturation and antibody production. A variety of major cardiac proteins, including cardiac myosin and troponin I, are targeted by auto-antibodies post MI (6). To assess binding of auto-antibodies to cardiac structures, CHF and control sera were used to stain frozen sections of healthy rat hearts. Strong staining was observed when using CHF serum, with a pattern resembling cardiomyocyte striations that confirmed auto-antibody binding to the cardiomyocyte contractile apparatus (Figure 3d).

**Figure 3:**
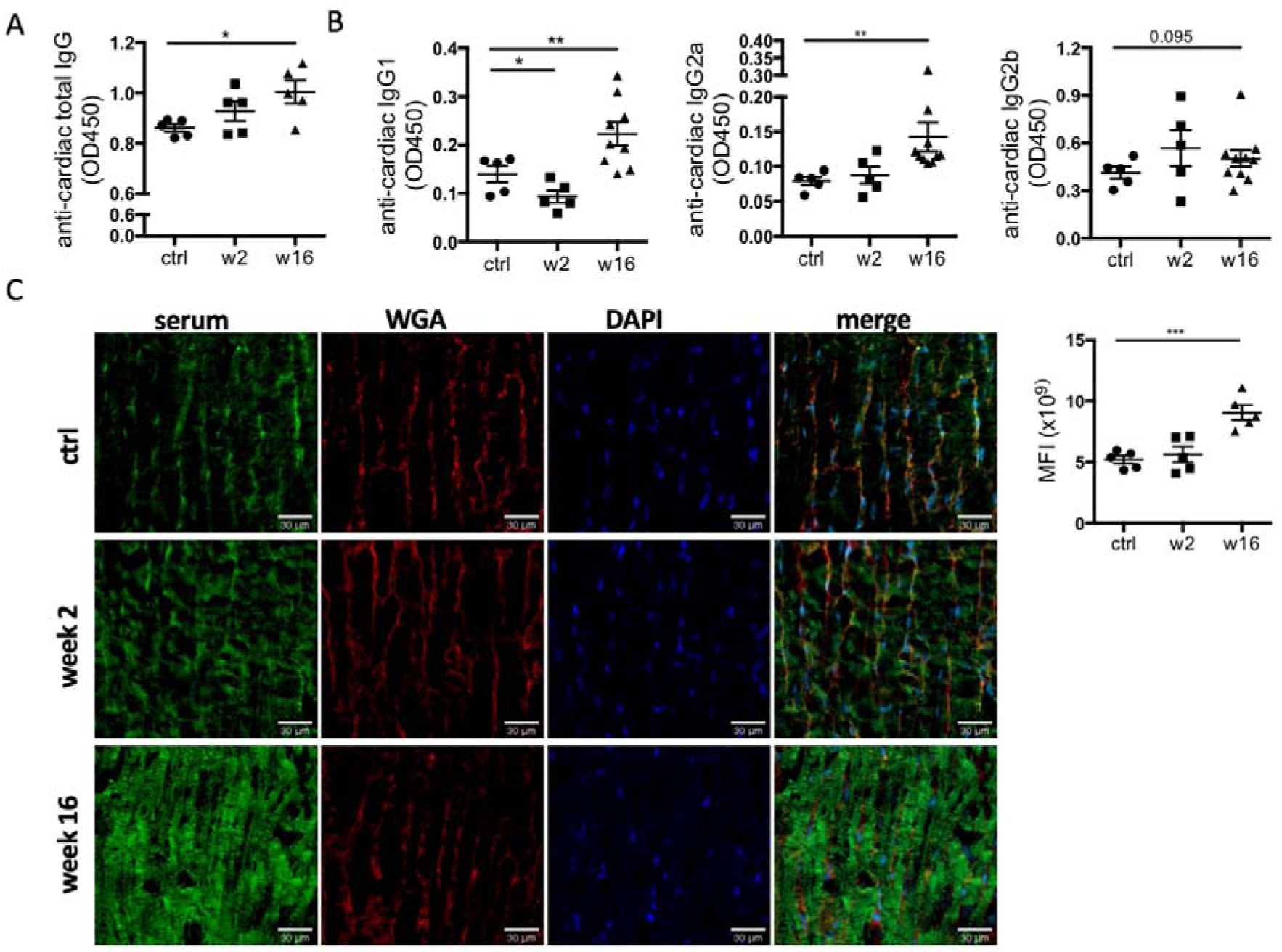
CHF serum contains anti-heart auto-reactive antibodies of mature class-switched isotypes. Myocardial infarction was induced by surgical ligation of the LAD. Serum was collected 2 weeks (acute post-MI) and 16 weeks (chronic heart failure stage) post-MI and analyzed by ELISA and immunofluorescence microscopy for the presence of heart specific auto-antibodies. A, B: ELISA using post-MI serum against rat heart lysate (protein fraction) showing a progressive increase post-MI in the levels of (A) total IgG (n = 5 / group) and (B) IgG1, IgG2a, and IgG2b isotype auto-antibodies reactive against the heart (n = 5 (ctrl, week 2) and 10 (week 16) / group). C: Immunofluorescence staining using post-MI serum on frozen sections of healthy rat hearts to assess binding of auto-antibodies to cardiac structures (n = 5 / group). A staining pattern resembling cardiomyocyte striations is observed indicating binding to the cardiomyocyte contractile apparatus. Overall staining intensity was quantified by measuring mean fluorescence intensity (MFI) using FIJI/ImageJ. Statistics: values represent mean +/- s.e.m., two-tailed unpaired Student’s t-test with Welch correction. *P<0.05, **P<0.005, ***P<0.001.

### 4) CHF serum creates a cytotoxic and hypertrophic environment for cardiomyocytes

The action of mature class-switched auto-antibodies has been implicated in immune-mediated tissue damage through complement- or cell-mediated cytotoxicity (15). To assess potential cell-independent pathological effects of CHF serum on cardiomyocytes, primary adult cardiomyocytes were isolated from healthy rats and treated with CHF and control sera for 24 hours. We observed significant cytotoxicity with CHF serum (Figure 4a), and an increase in numbers of caspase3+ apoptotic cells (Figure 4b, Supplementary Figure 2a). To investigate potential hypertrophic responses to CHF serum while avoiding changes in expression levels due to cytotoxicity, we used the robust cardiomyocyte cell line HL1-6, which is a defined homogenous sub-clone of HL-1 cells (16). To reduce baseline hypertrophy and ensure responsiveness to mild hypertrophic stimulation, HL1-6 were starved of epinephrine for two weeks before serum stimulation, and a decrease in cell size as measure of reduced baseline hypertrophy was confirmed (Supplementary Figure 3a). Studies investigating hypertrophy commonly use an increase in *Myh7, Nppa* and *Nppb* gene expression and a shift in the expression ratio between adult *Myh6* to embryonic *Myh7* as indicators of hypertrophy (17)(18)(19). The *Myh7*/*Myh6* ratio was indeed increased in cardiomyocytes after 48 hours of serum stimulation with 16 week post-MI CHF serum (Figure 4c) showing a mild but significant level of re-expression of embryonic *Myh7* indicating cardiomyocyte hypertrophy. *Nppa* and *Nppb* remained unchanged (Supplementary Figure 3b). Stimulating HL1-6 with 10μM of norepinephrine (NE) as hypertrophy positive control confirmed their ability to upregulate *Myh7* and *Nppb*, while *Nppa* remained unchanged (Supplementary Figure 3c) as suggested previously to be sufficient to induce hypertrophic growth (20)(21).

**Figure 4:**
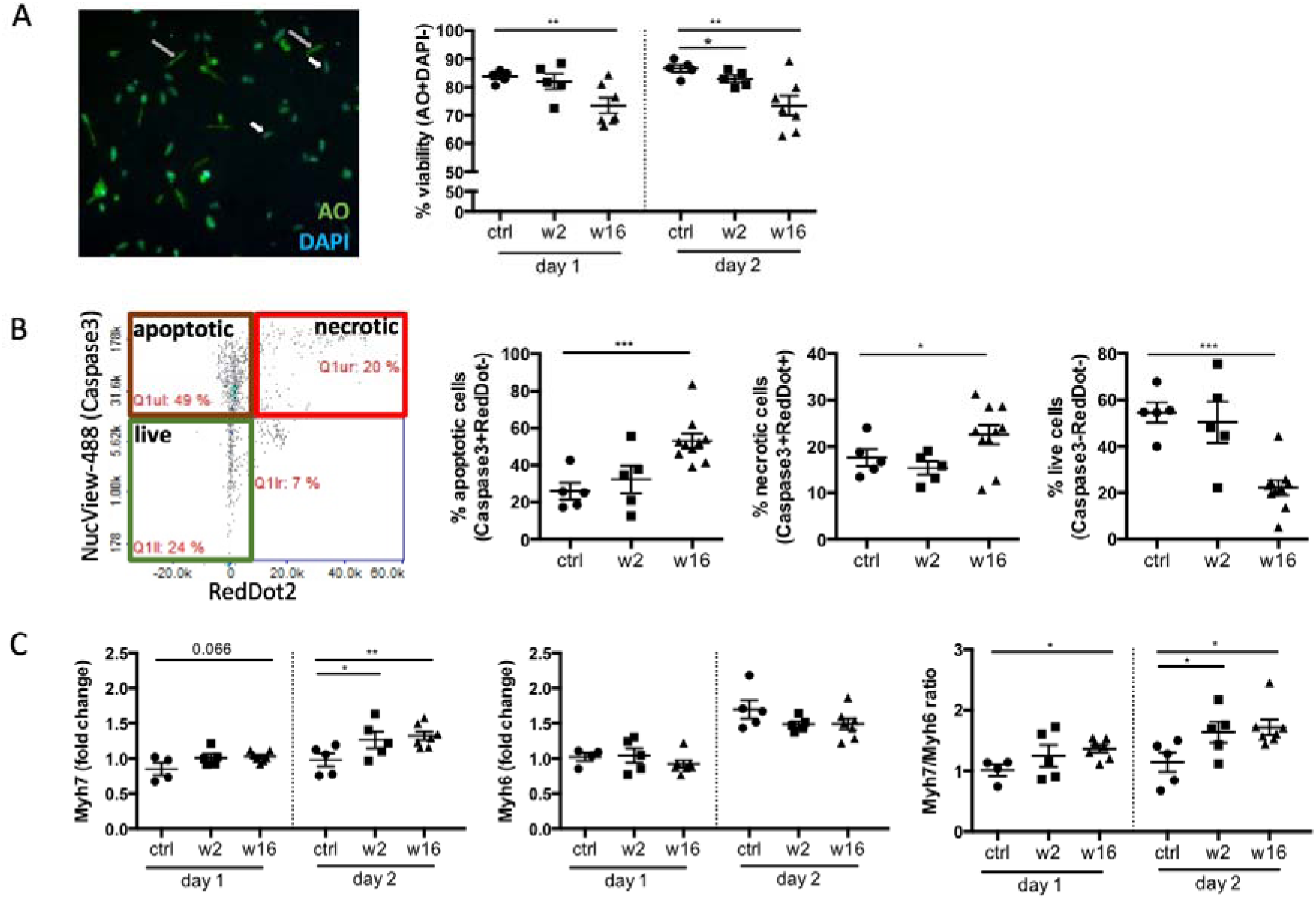
CHF serum creates a cytotoxic and hypertrophic environment for cardiomyocytes. Cardiomyocytes were isolated from healthy adult rats and treated with post- MI serum. Assays were performed using Via1-cassettes™ and Nucleocounter NC-200™ kindly provided by Chemometec, Denmark A: Viability assay staining cardiomyoctes with Acridine Orange (AO, total cell population) and DAPI (necrotic cell population), n = 5 (ctrl, week 2) and 7 (week 16). B: Cardiomyocyte apoptosis assay using NucView ™ Caspase 3 substrate (early apoptotic cells) and RedDot ™2 far red nuclear stain (necrotic cells). Images generated by the Nucleocounter NC-200™ were manually analyzes using FIJI image processing software, (n = 5 (ctrl, week 2) and 10 (week 16). C: qPCR assays testing the expression of Myh6 and Myh7 and beta-2 microglobulin as internal reference gene. Data analysis was performed using the 2^−ΔΔCt^ method to achieve a fold change over cells stimulated with serum from control rats, n = 4/5 (ctrl, week 2) and 8 (week 16). Statistics: values represent mean +/- s.e.m., two-tailed unpaired Student’s t-test with Welch correction. *P<0.05, **P<0.005, ***P<0.001.

## DISCUSSION

In this study, we document the presence of pathological adaptive immune auto-reactivity against the heart during CHF. Humoral factors in CHF serum directly damaged healthy cardiomyocytes *in vitro* by inducing apoptosis, decreasing cell survival and triggering a hypertrophic expression profile (**Figure 5**). We thus provide experimental evidence for the notion that aberrant activation of the adaptive immune system after MI promotes adverse ventricular remodeling leading to heart failure. Analysis of heart-draining mediastinal LN revealed a high number of IgG2a positive cells, representing mature B cells that class-switched to IgG2a.

**Figure 5:**
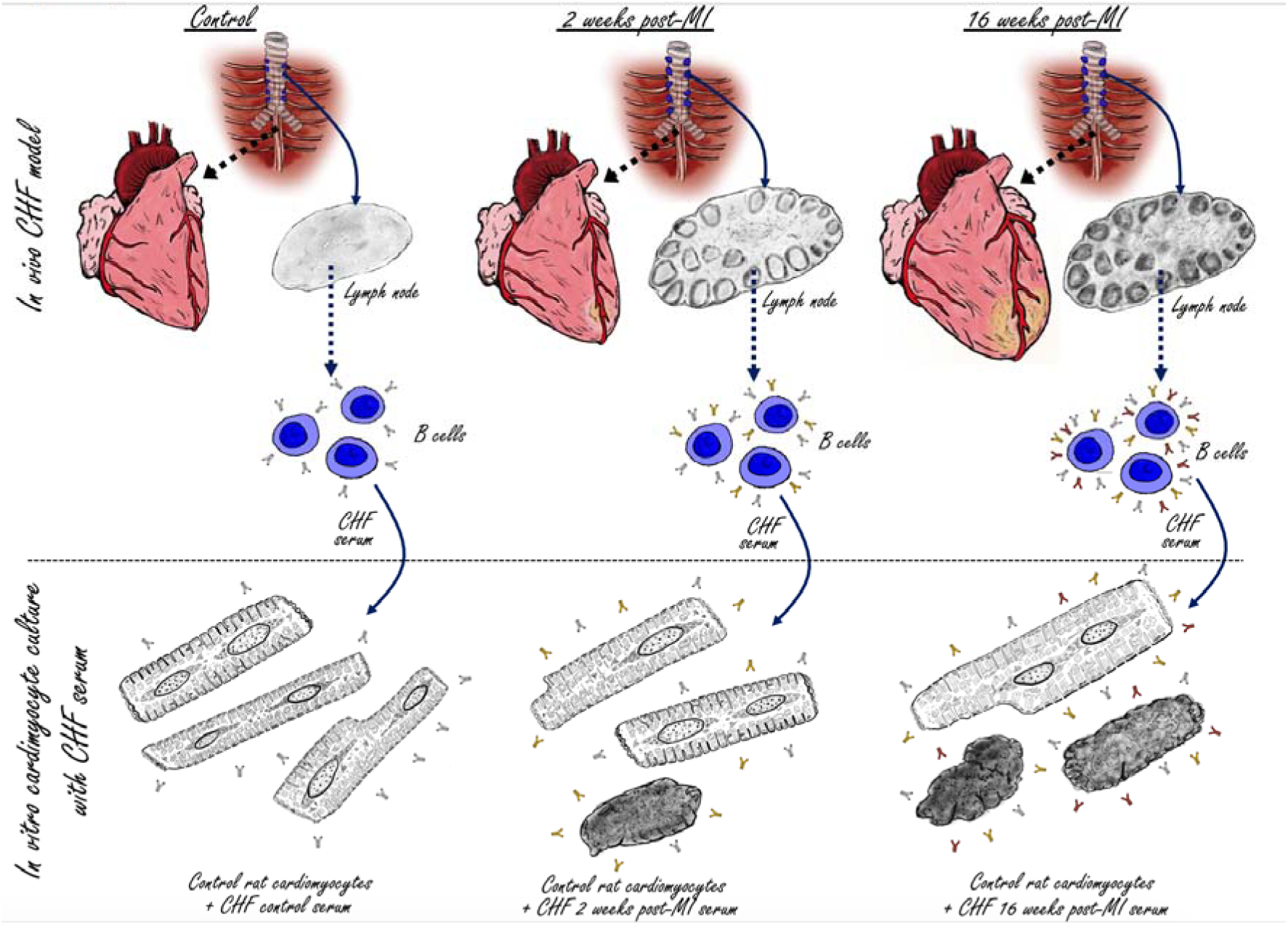
Visual abstract; the adaptive anti-heart immune response in CHF. A myocardial infarction initiates an adaptive immune response with activation of the heart draining mediastinal lymph nodes, formation of secondary follicles and B cell maturation with production of mature self-reactive auto-antibodies which induce cardiomyocyte cell death.

Upon activation, immature B cells undergo either clonal expansion for immediate antibody production or germinal center reactions in the lymph nodes. The germinal center reaction including somatic hyper-mutation and class-switch recombination, creates high affinity antibodies with variable Fc portions (22). The Fc portion of antibodies is of particular importance as it conveys their effector functions, which include (a) direct induction of cell death by receptor cross-linkage or blockade of receptor-ligand interactions, (b) recruitment of effector cells for antibody-dependent cell-mediated cytotoxicity (ADCC) or (c) antibody-dependent cellular phagocytosis (ADCP) by engagement of activating Fcγ receptors (FcγR) or complement-dependent cytotoxicity (CDC) (23). Depending on specificity and isotype, anti-cardiac auto-antibodies in CHF can therefore result in cardiac injury in a variety of ways (24). They mediate physiological damage by cross reacting with β1–adrenergic receptors, which play a role in heart contractility via sympathetic stimulation. The activation of β1–adrenergic receptors by auto-antibodies results in excessive stimulation which can induce left ventricular hypertrophy and pathological remodeling (25). Most notably, antibodies of IgG2 isotype including IgG2a and IgG2b play well established pathogenic role in autoimmune diseases such as systemic lupus erythematosus (SLE) (26). As we reported previously for an Resiquimod-induced SLE model, anti-cardiac antibodies of IgG2a and IgG2b isotype are present in circulation and deposited in the hearts (27). IgG auto-antibodies activate the complement cascade, which culminates in formation of the cytolytic membrane attack complex (MAC) comprising of complement C5b-9. The MAC forms a trans-membrane pore leading to necrotic death of the target cell (28). At sub-lytic concentrations however, MAC can induce caspase activation and apoptosis (29). Complement is accepted as a mediator of additional damage during acute MI and after reperfusion (30)(31), but more subtle sub-lytic activity has also been implicated in development of dilated cardiomyopathy. C5b-9 correlates with myocardial immunoglobulin deposition and expression of TNF-α (40). *In vitro*, C5b-9 attack on cardiomyocytes induces nuclear factor (NF)-κB activation as well as transcription, synthesis, and secretion of TNF-α by the cardiomyocytes themselves (32). NF-κB activation and TNF-α both induce cardiomyocyte hypertrophy (33)(34) *in vitro*, and the presence of immune cells able to respond to auto-antibody deposition *in vivo* will further increase the number of pathological pathways that contribute to CHF. In addition, as others have observed, an increase in *Myh7* upon CHF serum stimulation of HL1-6 cardiomyocytes, without corresponding increase of *Nppa* and *Nppb*, may still reflect detrimental effects (22). Notably, transgenic mice overexpressing *Myh7* had more progressive and severe cardiac damage than their wildtype counterparts (20), suggesting that an increase in *Myh7* alone is sufficient to cause cardiac deterioration. Additional cardiomyocyte responses induced by cell-free serum containing auto-antibodies, namely necrotic cell death, activation of caspase 3 as well as upregulation of *MyH7*, are likely mediated by direct complement-induced effects.

In summary, we show that long after the initially triggering MI, CHF serum still carries the ability to induce pathophysiological changes in healthy cardiomyocytes. This confirms that post-MI immune auto-reactivity is indeed a crucial contributing factor to adverse remodeling in remote areas of the myocardium, and offers new avenues for therapeutic intervention. Therapies may include immunomodulation to achieve an appropriate balance between inflammatory and regulatory immune cell populations and most importantly restore immunological tolerance to the heart. A large number of clinical trials have attempted to improve post-MI outcome by immunomodulation, with a strong focus so far on the innate immune system and short term readouts (35). The CANTOS trial using Canakinumab, a monoclonal antibody that neutralizes IL-1β, was the most recent attempt of immunomodulatory therapy aiming to prevent secondary infarcts in post-MI patients with elevated inflammatory profile (36). The beneficial effects of cardiomyocyte antigen-specific tolerogenic dendritic cells (DC) on post-MI function and remodelling in mice (37) provides a first proof of concept that post-MI tolerogenic immunotherapy targeting adaptive immunity may also be translatable into clinical use. Strikingly, several routine post-MI pharmacotherapies, including statin treatment, may also exert some of their beneficial effects by modulating the immune response. Statins in particular boost regulatory T-lymphocytes whilst inhibiting pro-inflammatory T-lymphocyte subpopulations (38). Importantly, identification of CHF as another member of the expanding family of immune-mediated diseases opens the door for new experimental, diagnostic and therapeutic approaches.

## ACKNOWLEDGEMENTS

We are grateful to members of Prof Harding’s and Prof Gorelik’s groups for technical support and helpful discussions, to Prof Michael Schneider, Prof Nick Peters and Dr Rasheda Chowdhury for providing access to essential equipment and for sharing reagents. We would also like to thank the team at Chemometec for their support in generating the data with the NucleoCounter NC-200 and NC-3000 automated cell counters. We thank staff at the animal facility at Imperial College London for help with animal husbandry and maintenance. The authors further acknowledge the use of the Facility for Imaging and Light Microscopy (FILM) at Imperial College London. We are grateful to Dr Rosalinda Doty, Director of Pathology Services, The Jackson Laboratories, for advice on lymph node histopathology.

## AUTHOR CONTRIBUTIONS

SS conceived and designed the study. SS, AS, CM, EF, SeR, PS, SR, SN, KS and JLSA designed, planned, performed and analyzed experiments. SS and AS wrote the manuscript. SEH, NR, JG and SS provided financial support. MH, SHE, JG and NR advised and revised the manuscript.

## MATERIALS AND METHODS

### MI surgery

All animal procedures were approved by the Imperial College Governance Board for Animal Research and in accordance with the UK Home Office Animals (Scientific Procedures) Act 1986 and Directive 2010/63/EU of the European Parliament on the protection of animals used for scientific purposes. 30 male Sprague Dawley rats (250 to 350 g) and10 male Lewis rats (250 to 350 g) were obtained from Charles River Laboratories, UK. They were fed standard rat chow ad libitum. Rats were housed at a density of 4-5 per cage and maintained on a 12-hour light/dark cycle at 21°C. MI was induced by LAD ligation following a previously described procedure (39). Briefly, anesthesia was induced in an induction chamber with 5% isoflurane. Rats were then intubated, and anesthesia was maintained at 2% isoflurane. Perioperative analgesic regime included buprenorphine (0.05 mg/kg) and Rimadyl (5 mg/kg), Baytril (5 mg/kg) for prophylaxis, and 1 ml of 0.9% saline administered sub-cutaneously to ensure appropriate hydration. The heart was visualized by a left parasternal incision followed by thoracotomy through the fourth intercostal space and removal of the pericardium. Ligation of the LAD was performed 1-2 mm distal to the inferior border of the left atrium using an 7-0 Prolene suture. Blanching and cyanosis of the left ventricular free wall and apex were used as confirmation of efficient ligation. The thoracotomy and chest wall were closed using 4-0 coated Vicryl sutures, and the skin was closed using 4-0 Vicryl sutures. Daily doses of Rimadyl (5 mg/kg) were administered for at least 48 hours after surgery for post-operative analgesia.

### Adult rat cardiomyocyte isolation

Adult rat cardiomyocytes were isolated as described previously (40). Briefly, hearts were excised and placed in ice cold Krebs-Henseleit (KH) Buffer (119mM NaCl, 4.7mM KCl, 0.94mM MgSO4, 1mM CaCl2, 1.2mM KH2PO4, 25mM NaHCO3, 11.5mM glucose; 95% O2, 5% CO2), cannulated via the aorta and perfused with KH at 37°C using a Langendorff apparatus. When blood was cleared from the coronary circulation, the KH buffer was switched to a low calcium (LoCa2+) buffer (12-15μM CaCl2, 120mM NaCl, 5.4mM KCl, 5mM MgSO4, 5mM pyruvate, 20mM glucose, 20mM taurine, 10mM HEPES, 5mM nitrilotriacetic acid (NTA); 100% O2) to stop contraction. A solution of 1mg/ml Collagenase II and 0.6mg/ml Hyluronidase (C+H) in enzyme buffer (12-15μM CaCl2, 120mM NaCl, 5.4mM KCl, 5mM MgSO4, 5mM pyruvate, 20mM glucose, 20mM taurine, 10mM HEPES,150μM Ca2+) was then perfused into the heart for 10 minutes after which the heart was minced in fresh C+H buffer. Minced samples were shaken mechanically at 35°C for 5 minutes and the supernatant was filtered through gauze and fresh C+H replaced in the tube with undigested tissue. The samples were shaken again for a further 30 minutes and the supernatant was strained through gauze. The resulting filtrates were centrifuged for 1 minute at 700rpm which formed a pellet of isolated cardiomyocytes. Supernatants were removed from these samples and the pellets re-suspended in enzyme buffer. This method yields cardiomyocytes which are tolerant of Ca2+, quiescent when not stimulated and can be maintained in cell culture free of contamination for 4 days. Isolated primary adult cardiomyocytes were cultured in modified M199 medium (Thermo Fisher Scientific, Rochford, UK) containing bovine serum albumin (0.5 g/L), creatine (5 mmol/L), taurine (5 mmol/L), L-ascorbic acid (100 μmol/L), carnitine (2 mmol/L), and penicillin/streptomycin (100 mmol/L) and treated with post-MI sera at a dilution of 1:10 for 24h and 48h.

### HL-1 cell culture and stimulation

HL-1-6 cells were kindly provided by Dr Emauel Dupont, Imperial College London, and cultured as previously described in 5% CO2 at 37°C in Claycomb medium (Sigma-Aldrich, Gillingham, UK) supplemented with 10% fetal calf serum (Biosera, France), 1% L-Glutamine and 100 μM norepinephrine (both Sigma-Aldrich, Gillingham, UK) (16). Complete Claycomb medium was changed every 48 hours and cells were passaged using 0.05% Trypsin-EDTA and Trypsin inhibitor, soybean (both Sigma-Aldrich, Gillingham, UK). Before serum stimulation, cells were starved of NE for 2 weeks. Cells were stimulated with post-MI serum diluted 1:100 in culture medium for 24 and 48h. Positive controls were supplied with fresh NE.

### qPCR

After discarding cell-culture medium, RNAlater was added for 1 minute and cells were scratched off the surface using TRIzolTM (both Gibco, Thermo Fisher Scientific, Dartford, UK). RNA isolation was performed using the phenol-chloroform extraction method. Briefly, after defrosting cells in TRIzolTM, chloroform (Sigma-Aldrich, Gillingham UK) was added and mixed thoroughly by vortexing for 15 seconds. The samples were incubated for 15 minutes on ice then centrifuged for 15 minutes at 12,000rcf, 4**°**C. The top aqueous phase was transferred to new microcentrifuge tubes. Equal volumes of isopropanol (Fisher Scientific, Loughborough, UK) were added and samples were incubated on ice for 10 minutes followed by centrifugation for 10 minutes at 12,000 rcf, 4**°**C. The pellet was washed in 70% Ethanol, left to air dry and resuspended in 25ul of DNase/RNase-free distilled water (Invitrogen, Thermo Fisher Scientific, Rochford, UK). RNA concentration and purity was determined by using a NanoDrop spectrophotometer (ND-8000-GL, Thermo Fisher Scientific, Rochford, UK). RNA concentration was adjusted to 50ng/μl and cDNA synthesis was performed using the High capacity cDNA Reverse Transcription kit (Applied Biosystems, Thermo Fisher Scientific, Warrington, UK) according to the manufacturer’s instructions. The reaction was carried out using the thermal cycler GeneAmp® PCR System 2700 (Applied Biosystems, Thermofisher Scientific, Warrington, UK). qPCR was performed using the Eppendorf Mastercycler® RealPlex2, TaqMan® Gene Expression Master Mix (Thermo Fisher Scientific, Rochford, UK), and TaqMan® Gene Expression Assays following manufacturer’s instructions. For MCEC-1 experiments, Vcam1 (Mm01320970_m1), Icam1 (Mm00516023_m1) and E-selectin (Mm00441278_m1) expression assays was used. For HL-1 experiments, Myh7 (Mm00600555_m1), Myh6 (Mm00440359_m1), Nppa (Mm01255747_g1), Nppb (Mm01255770_g1) was used. Beta-2 microglobulin (Mm00437762_m1) served as internal reference gene. Threshold cycle (C_t_) values of the target genes were normalized to the experimental control. Data analysis was performed using the 2^−ΔΔCt^ method.

### Immunofluorescence staining

For analysis of **auto-antibody binding**, healthy rat hearts were embedded and frozen in OCT medium (Sigma-Aldrich, Gillingham, UK) and cut into 5μm sections. Control, 2 and 16 weeks post-MI sera were used as primary antibodies in 1:100 dilution in PBS and incubated overnight at 4°C. Sections were washed three times with PBS and a secondary anti-rat IgG AlexaFluor488 antibody (BioLegend, London, UK) was used for detection. Wheat germ agglutinin (WGA)-AlexaFluor488 (Invitrogen, Thermo Fisher Scientific, Rochford, UK) was used as membrane counterstain. Sections were mounted with hard setting mounting medium containing DAPI (Vectashield, Vector Laboratories, Peterborough, UK). For detection **of IgG2a+ cells in mediastinal lymph nodes**, frozen LN sections were stained with Alexa Fluor® 488 Goat anti-rat IgG and Alexa Fluor® 647 anti-rat IgG2a (both BioLegend, London, UK). Sections were mounted using DAPI-containing mounting medium (Abcam, Cambridge, UK). For **HL-1-6 cell-size analysis**, HL-1 cells were seeded into chamber slides, fixed with 4%PFA for 15 minutes and incubated with WGA for 15 minutes. Slides were mounted with DAPI containing mounting medium (Abcam, Cambridge, UK). Images were captured using a LMD7000 microscope (Leica microsystems, Milton Keynes, UK) or a Zeiss Axio Observer inverted microscope and processed using the public domain software ImageJ (NIH; http://rsb.info.nih.gov)(41).

### ELISA

The ELISA protocol for **detection of rat anti-heart auto-antibodies** was optimized using mouse, rat, pig lysate and titration of serum concentration to achieve the best possible background to signal ratio (Supplementary Figure 4). The final standard protocol used 4*μ*g/*μ*l pig heart lysate as capture reagent and sample serum dilutions of 1:10 and 1:100 in PBS. ELISAs plates (SpectraMax Paradigm Molecular-Devices, UK) were coated with 50μl per well of 4μg/μl pig heart lysate lysate (Novus Biologicals, Bio-Techne, Abingdon, UK) diluted in PBS overnight at 4°C. Plates were washed three times for 5 minutes each with 200μl per well of ELISA washing buffer (0.1% Tween 20 in PBS). Then, 50μl of the rat sample sera per well was added in either 1:10, 1:100 dilutions for overnight incubation at 4°C. Detection reagents were either anti-rat IgG-HRP, anti-rat IgG1/2a/2b/2c-Biotin combined with Streptavidin-HRP Streptavidin-HRP (all BioLegend, London, UK).

For relative **quantification of antibody isotypes** in the serum, a Rapid Antibody Isotyping Assay Kit, rat (Thermo Fisher Scientific, Rochford, UK) was used according to the manufacturer’s instructions using previously determined serum dilutions (42). For all ELISA assays, detection steps were performed using a TMB ELISA buffer kit (Peprotech, London, UK) as per manufacturer’s instructions. An ELISA plate reader (SpectraMax Paradigm Molecular-Devices, UK) was used to measure light absorbance at 450nm for the HRP product and 570nm for the plastic background.

### Cardiomyocyte viability and cell cycle assay

Adult primary cardiomyocytes were harvested on days 1 and 2 post-culture and analyzed using the image cytometry system Nucleocounter NC-200 and NC-3000 (Chemometec, Denmark) and Via1-Cassette(tm)as per manufacturer’s instructions. NucleoCounter counts cells using. Briefly, Via1-Cassette(tm) stains all cells with acridine orange (AO) and dead cells with DAPI. Apoptosis assays were performed using Nucleocounter NC-3000 and a Flexicyte apoptosis/necrosis detection kit based on staining with caspase 3 substrate NucView 488 and RedDot 2 (all Biotium, Cambridge Bioscience, UK). Apoptotic cells are defined as NucView488 positive and RedDot2 negative, necrotic cells are defined as NucView488 positive and RedDot2 positive (Supplementary Figure 2a). Images obtained or all assays were analyzed manually using the FIJI (ImageJ) to quantify percent viability. DAPI staining intensity representing individual cell cycle stages due to differences in DNA content was measured for cell cycle analysis (Supplementary Figure 2a).

### Experimental design and statistical analysis

Animal number and sample sizes calculations were performed using G*Power 3.1 (43) available at http://www.gpower.hhu.de/ and reflect effect sizes obtained in previous experiments with comparable readouts. Statistical analysis was performed using SPSS or GraphPad Prism. Normally distributed data were presented as mean±s.e.m., and one-or two-tailed unpaired or paired Student’s t-tests were performed as appropriate for parametric data and indicated in the respective figure legends. Healthy controls and CHF groups are assumed to have different standard deviations due to variation in MI severity, thus Welch’s correction was applied. Exact p-values, t-values and degrees of freedom for each experiment are provided in the supplemental information documents.

## MATERIALS & CORRESPONDENCE

Please address correspondence to Dr Susanne Sattler.

## FIGURES AND FIGURE LEGENDS

**Supplementary Figure 1.**
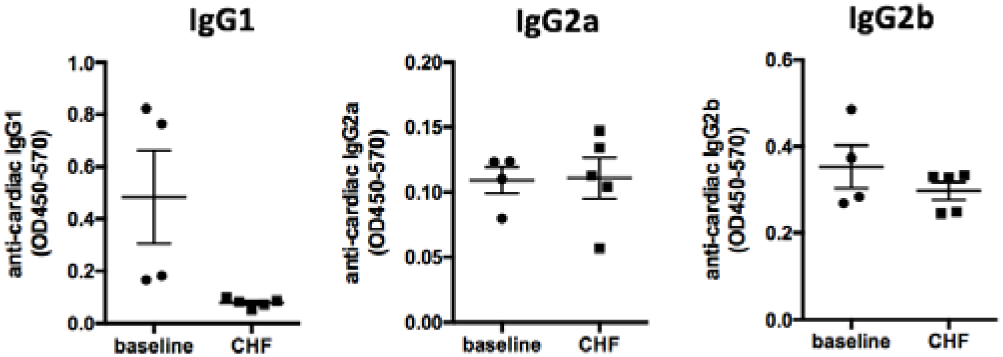
Anti-heart autoantibodies in Lewis rats with CHF. Myocardial infarction was induced by surgical ligation of the LAD. CHF serum was collected 16 weeks post-MI and analyzed by ELISA against full heart lysate for the detection of heart specific auto-antibodies. IgG1, IgG2a and IgG2b isotype autoantibodies are not increased over baseline during CHF in Lewis mice. Statistics: n = 4 (ctrl) and 5 (CHF), values represent mean +/- s.e.m., two-tailed unpaired Student’s t-test with Welch correction.

**Supplementary Figure 2.**
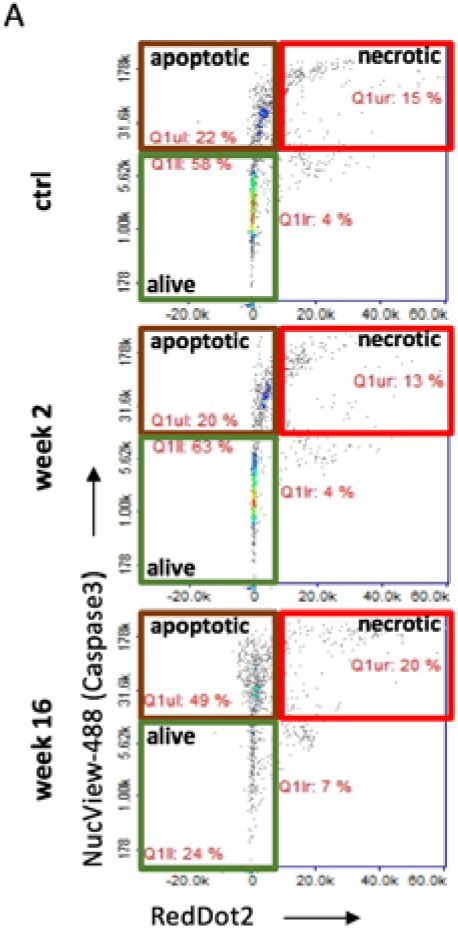
Apoptosis assay. Cardiomyocytes were isolated from healthy adult rats and treated with post-MI serum. Assays were performed using Via1-cassettes™ and Nucleocounter NC-200™ kindly provided by Chemometec, Denmark. Example plots obtained from the cardiomyocyte apoptosis assay using NucView ™ Caspase 3 substrate (early apoptotic cells) and RedDot™2 far red nuclear stain (necrotic cells).

**Supplementary Figure 3:**
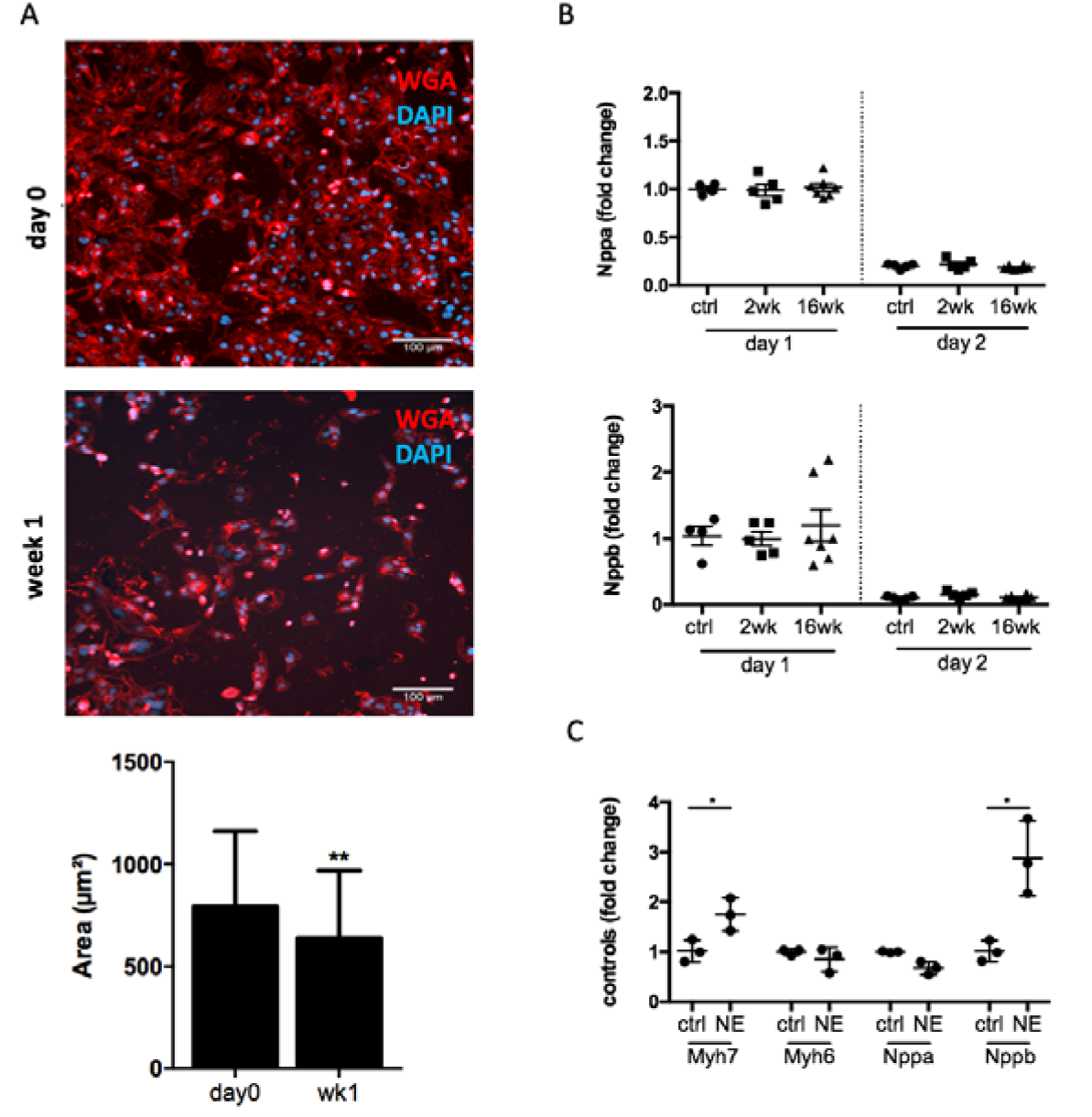
Characterization of HL1-6 cardiomyocyte line to assess suitability for hypertrophy experiments. A: Immunofluorescence staining with WGA (cell membrane) and DAPI (nuclei) and quantification of cell size (hypertrophy) of HL1-6 cells at baseline and after starvation off nor-epinephrine (NE) for 1 week. Statistics: values represent mean area +/-s.d. of >60 cells, two-tailed unpaired Student’s t-test. B: Cardiomyocytes were isolated from healthy adult rats and treated with post-MI serum. qPCR assays testing the expression of *Nppa* and *Nppb*. and beta-2 microglobulin as internal reference gene. Data analysis was performed using the 2^−ΔΔCt^ method to achieve a fold change over cells stimulated with serum from control rats, n = 4/5 (ctrl, week 2) and 7 (week 16). Values represent mean +/- s.e.m., one-tailed unpaired Student’s t-test with Welch correction. C: qPCR assays testing the expression of *Myh6, Myh7, Nppa* and *Nppb* in response to stimulation of HL1-6 cells with 10um NE used as positive control to assess HL1-6 response to a hypertrophic stimulus. Data analysis was performed using the 2^−ΔΔCt^ method using beta-2 microglobulin as internal reference gene to achieve a fold change over control, n=3/group, technical triplicates / group, values represent mean +/-s.d., two-tailed unpaired Student’s t-test. *P<0.05, **P<0.005.

**Supplementary Figure 4.**
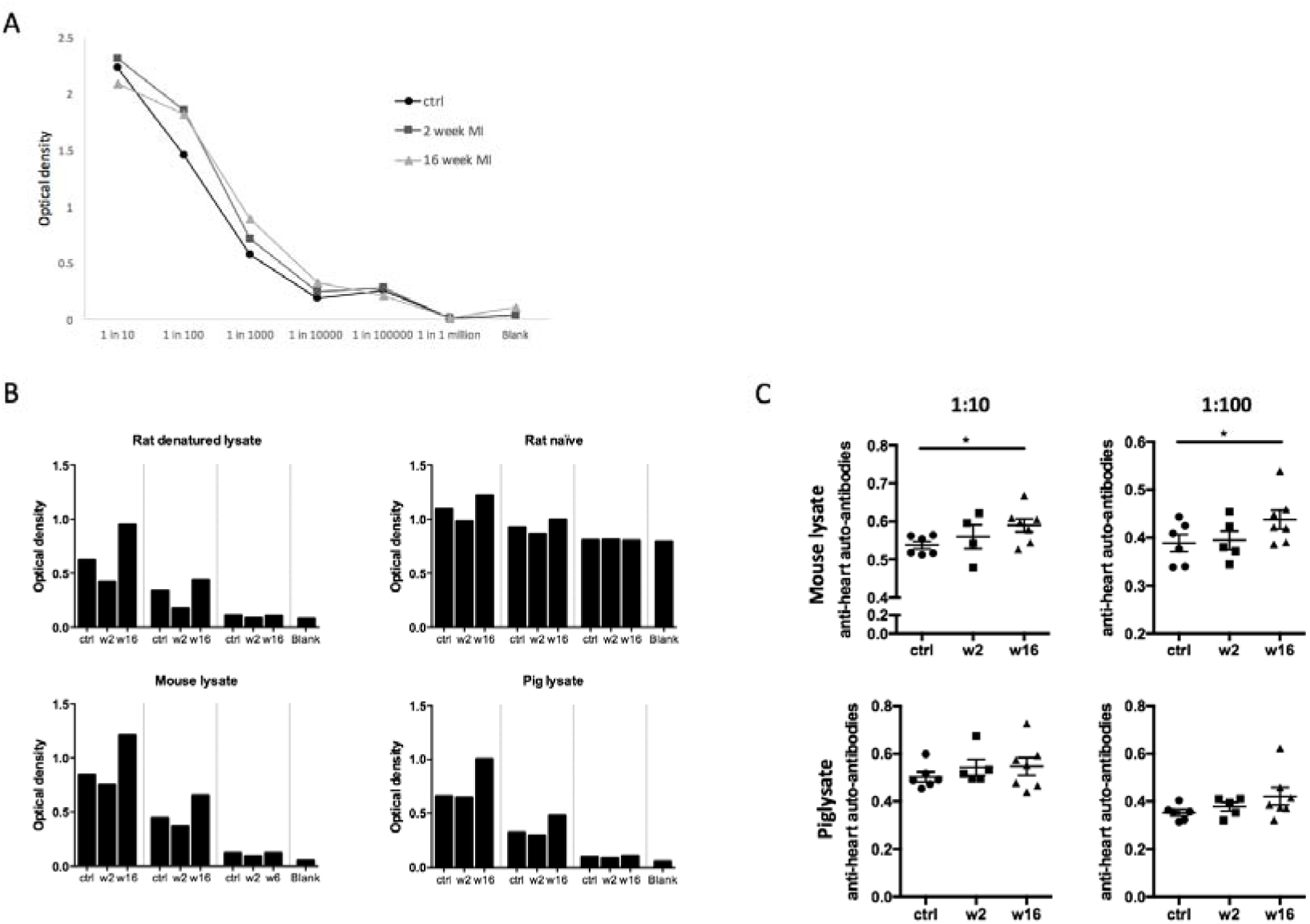
Setting up and optimization of anti-heart lysate ELISA. A: Serial dilutions of control, 2 week MI and 16 week MI rat sera (1:10 to 1:1 million). B: ELISA results for different lysates (rat denatured, rat naïve, mouse naïve, pig naïve) per control, 2 week MI and 16 week MI serum samples in 1:10, 1:100, 1:1000 dilutions. C: Repeat of ELISA assays coated with mouse and pig lysate for two different sample concentrations (dilutions of 1:10 and 1:100) for control (n=6), 2 week MI (n=5) and 16 week MI (n=7) serum samples. All experiments performed in technical triplicates. Statistics: n = 5-7 / group, values represent mean +/- s.e.m., one-tailed unpaired Student’s t-test with Welch correction. *P<0.05. Statistics: (A, B) n = 3-8 / group, values represent mean +/- s.e.m., two-tailed unpaired Student’s t-test with Welch correction. *P<0.05.

